# Additive and substitutable prey responses to feral and native predator combinations

**DOI:** 10.1101/2024.09.09.612031

**Authors:** Justin R. Saint Juliana, S.S. Bleicher, S. Mukherjee, V. Sundararaj, J.S. Brown, B.P Kotler

## Abstract

In captive experiments of Negev Desert granivores, we investigated the ways in which combinations of feral mesocarnivores and native predators alter wild prey behavior. We hypothesized that feral mesocarnivores would have a greater impact on prey energy acquisition, reflected in foraging dynamics, than native predators. Allenby’s gerbils (*Gerbillus andersoni allenbyi*) and greater Egyptian gerbils (*Gerbillus pyramidum*) were used as prey species, with feral dogs (*Canis lupus familiaris*), feral cats (*Felis catus*), barn owls (*Tyto alba*), and horned vipers (*Cerastes gasperetti*) as predators. Gerbil perceived risk was measured using optimal patch-use theory, with exposures to tethered predators occurring hourly throughout the night. Some nights, two predators were alternated every other hour. We found that human-commensal predators, particularly feral cats, induced stronger foraging than native predators, such as barn owls. Combined predators caused gerbils to decrease foraging only when a higher-risk predator was introduced, as indicated by higher giving-up densities (GUDs) for the dog and cat combination compared to the dog alone, and a nonsignificant increase compared to the cat alone. The impact of feral cats especially appears to outweigh that of native predators. This highlights the conservation challenges to arid environments where feral cats have become ubiquitous.

**Highlights:**

- Feral predators, particularly cats and dogs, elicit stronger anti-predator responses in Negev Desert gerbils than native predators, significantly altering foraging behavior and habitat use.
- The presence of feral predators in the Negev Desert leads to cumulative impacts on prey species, resulting in increased vigilance and reduced resource acquisition, which suggests a disruption of native predator-prey dynamics.
- The study’s findings emphasize the unique conservation challenges posed by human-commensal species in arid ecosystems, where limited refugia increases the predation risk from invasive and feral species.
- This research demonstrates how predator-prey interactions in the arid communities tested are impacted by both additive predator effects between native predators, and between feral predators, but when combining feral predators with native ones, the effect is substitutable with feral predators disproportionately influencing prey behavior.

**Graphical Abstract:** 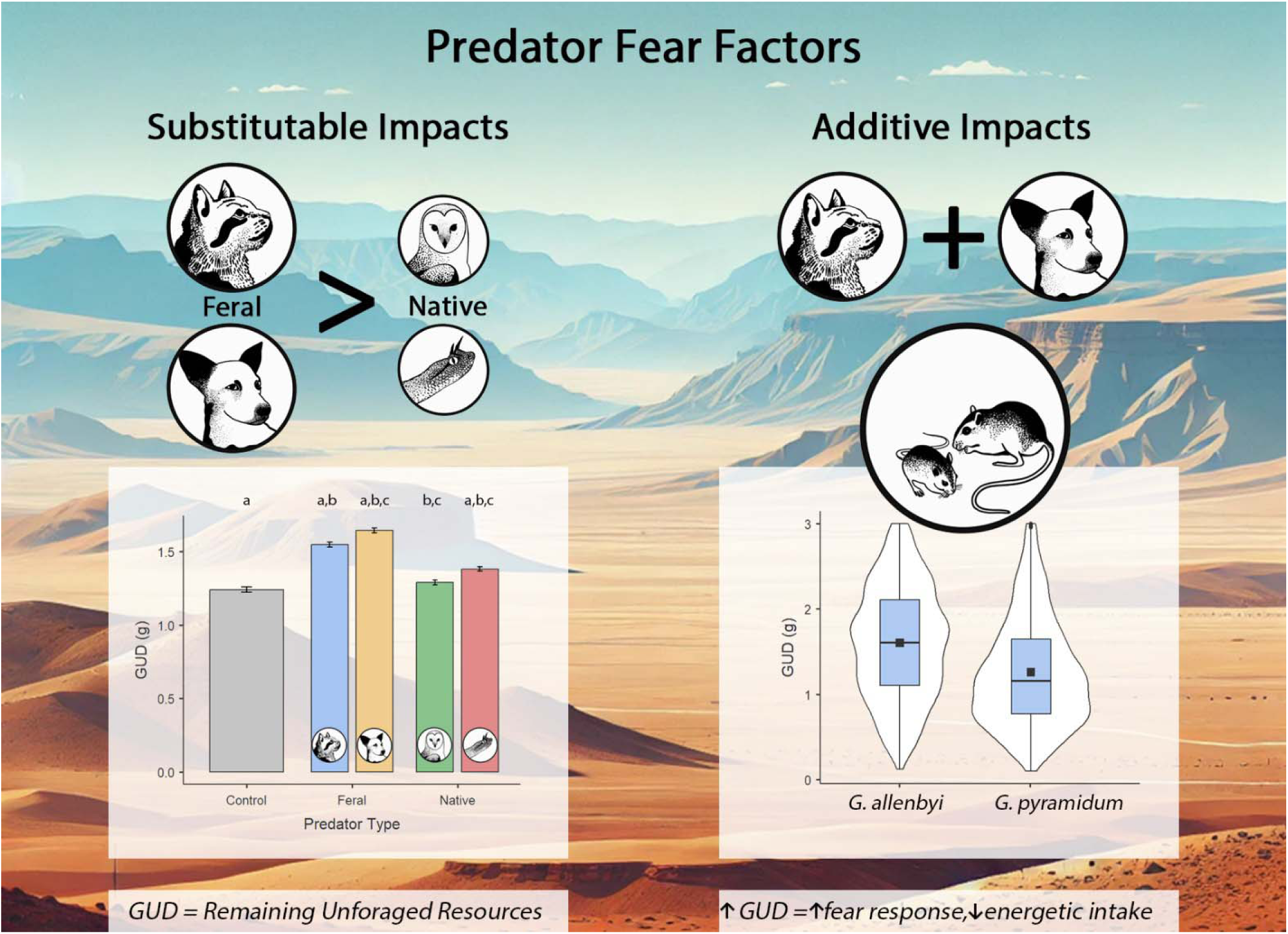

## Introduction

Predators are known to impact the richness of biological communities through top-down trophic interactions [1–3]. Predator-prey interactions enrich biological communities by allowing adaptive strategies of varying prey populations to coexist in space and time [4,5]. While the original perception of predator influence on communities was predominantly consumptive [6], the ecological research community has come to accept the non-consumptive impacts of predators to be equal or greater to those of direct predation [7–9]. Overall, increases in risk, and increases in the diversity of predators promote increased diversity of anti-predator adaptations among prey, which in turn help facilitate prey coexistence [10–13]. In a period of great biodiversity loss, especially predator loss [14], the impacts of changes in predator communities are expected to reverberate down through the prey communities. While the composition of predator assemblages has impacts globally, a disproportionate impact to predation risk has always been attributed in arid ecosystems, namely due to scarcity of refuge, despite a rich biodiversity of vulnerable prey species [15–17]. In addition, the studies evaluating the structure of predator assemblages have scarcely incorporated human-commensal, feral and invasive predators as part of the same analysis.

Despite the agreement on the impacts of risk on prey diversity, measuring risk intensity in empirical experiments has been proven complicated, and quantifying prey response to said variation is harder still. The current standard for testing impact of predation risk on prey, involves captive experiments, managing titration of food and safety through the addition of predators [18], or regulating available resources in the environment [19]. Current evidence that the effects of predator abundance on prey behavior tends to be cumulative, but non-additive [20–22]. For example, gerbils exposed to one owl decreased foraging by 18%, however the addition of a second owl only decreased foraging by 5% further and a third owl reduced foraging only by 1% [23]. This may be due to a “buy one get one half off” type of situation where the value of vigilance is greater when risk is higher. In other words, prey being vigilant for one owl, will also detect a 2^nd^ or 3^rd^ one. This type of benefit may apply to similar predators, but not predators that are very different. In contrast, with some predators, for example foxes, desert rodents tend to only “tune in” when the level reaches a certain threshold[24]..

While titration of risk is a clear influencer of risk accumulation, rarely is it ecologically relevant to observe a single predator species as the only source of risk in an environment. Multi-predator impacts have been predominantly studies in consumptive studies in aquatic and arachnid model communities, and have shown to take two major forms: facilitation or interference between the two predators [25,26].

Behaviorally, the impact of predators on each other depends on how prey perceive risk from these predators, thus impacting their habitat choices and the amount of time they spend foraging. Predator facilitation benefits predators in two major ways: 1. One predator drives prey towards the habitat preferred by another predator [25,27,28] in a mutualistic benefit. 2. Presence of one predator overpowers the attention given to another predator thus making the prey more vulnerable to the other predator that is less focused on, *i.e.*, a commensal relationship between the predators [29]. For many prey species, the risk from one predator requires much greater vigilance, or other types of defensive behavior, and thus marginalizes the attention (vigilance behavior) and energy expenditure (as decerned through time foraging) that can be allocated to avoidance of the lower risk predator [30,31]. It is important to be careful when interpreting the latter case when researching predation based on prey behavior. The predator that has a lower impact on the prey’s fitness may seem to cause no risk, because it is masked by the predator that the prey perceives to be riskier.

Predator facilitation should narrow the availability of perceived safe habitat and narrow the realized niche that the prey can occupy. This logically changes both community interactions, and the populations dynamics of both predators and prey. The lower the abundance and distribution of prey, the less fit both predators become in the long run. For example, the desert pocket mouse (*Cheatodipus penicillatus*), from the Mojave Desert, favors a strategy of torpor and reduction of foraging to a minimal sustainable level in response to combinations of owls and snakes in their environment [32]. When this occurs, neither predator is able to hunt effectively, despite the scenario suggesting facilitation. On the flip side, when predators interfere with each other, the prey should effectively be able to use more resources as a result of the conflict between the predators, reducing the overall risk [33,34]. This interaction is typically observed when the two predators use the same habitat, and drive the prey to use habitat that is less optimal for both predators. For example, when a population of gerbils is exposed to both owls and foxes, they prefer to forage under the safety of bushes where neither predator has easy access to them [35].

Different prey species will respond to the interaction of predators in different ways. This is a consequence of evolutionary changes driven both by predator avoidance [36,37] and by intraguild competition strategies pushing prey populations to evolve strategies that balance both competition and predation costs [30–32,38–41]. Specifically, the foraging strategy of competing prey species will alter how susceptible they may be to predation risk [42,43]. Examples include locomotion type, with saltatory motion conveying advantages for predator avoidance [44,45], sensory prowess as in the example of inner ear bullae allowing for extra sensitive hearing and early escape [46], and even unique special adaptations such as torpor as in the example of the abovementioned pocket mice [32]. In this experiment, we test the responses of two competing gerbil species from the Negev Desert in Israel, *Gerbillus andersoni allenbyi* (GA) and *Gerbillus pyramidum* (GP). We expect that the same experiments would result in different outcomes from species applying different anti-predator strategies. Specifically, the two gerbil species differ in their size with GA being significantly smaller than GP [47]. The size difference between those two well-studied model species has given rise to significant behavioral differences both in intra specific competition strategies and management of predation risk. The variation between the species, interspecific competitive dynamics between them, and intraspecific competition within the populations combined with the richness of the environment should also alter the impact of combined predators. We expect the addition of predators to have a greater impact when the competition for resources is lower (*cf.* Makin and Kotler 2020).

This study aims to compare the response of two desert gerbil species to titrations of predation risk through the exposure to four different predators: two human-commensal species (a feral cat and a dog), a barn owl, and an olfactory cue of a horned viper in the form of musk imbued sand. We intend to test each predator alone and in combination with each of the other predators to assess their cumulative and additive impacts on prey behavior, or whether their impacts are substitutable.

Given excessive impacts of human-commensal predators on small mammal communities worldwide [49,50] we predict a potentially disproportionate assessment of the risk these predators pose compared with native predators. The directionality of disproportionality may increase perceived risk, if the prey species understand the risks they pose, but may just as likely be significantly lower if the prey are naïve to the risks they pose [51]. Despite expected differences between the prey species’ behavior, we predict that the general response to predator stressors will follow similar patterns, as their anti-predator evasion strategies are similar enough. We predicted that the combination of any two predators would result in additive responses of the behavior of the gerbils. Additionally, we predicted that the closer the hunting strategy of any two predators, the lower the added impact will be on gerbil apprehension and foraging. *i.e.* adjustment in behavior applied towards one predator would automatically apply to the similar predator. For example, the response of gerbils to a fox would be to move the majority of their foraging under the safety of bushes. The logical response to a jackal in the environment would be very similar, therefore the response would not vary much when both are present. However, when avoiding snakes that ambush under bush cover, foraging in the open is the logical safe response [33,48]. Therefore, combing a fox and a snake would increase the risk throughout the environment, thus registering as a much greater threat than two ambush predators, or two active searching predators. (Table 1). Where the risk from one predator type outweighs the risk from another, the effect may register as overshadowing of the impact of the lower-risk predator [30].

**Table 1.** Predictive patterns of gerbil behavior in relation to four predators used in the experiment.

| **Predator** | **Safer**<br>**microhabitat**<br>**for gerbils** | **Predator**<br>**hunting**<br>**activity type** | **Potential**<br>**interference** | **Expected**<br>**facilitation** | **Potential**<br>**replacement**<br>**to** |
| --- | --- | --- | --- | --- | --- |
| Cat | Bush | Ground<br>foraging | Dog, Owl | Snake | Snake |
| Dog | Bush | Ground<br>foraging | Cat, Owl | Snake | Snake |
| Snake | Open | Ambush |  | Cat, Dog, Owl |  |
| Owl | Bush | Aerial | Cat, Dog | Snake | Snake |

## Methods

### Species

For this experiment we used two common gerbils from the Negev Desert of Israel: the greater Egyptian gerbil *Gerbillus pyramidum* (GP), 40 g, and Allenby’s gerbil *Gerbillus andersoni allenbyi* (GA), 24 g [52]. Both these gerbils are nocturnal, are commonly found on sandy substrates such as sand dunes, and are considered granivores, despite foraging on available insects, and green vegetation when available [53]. Both gerbils have adaptations to reduce the risk of predation, including saltatorial locomotion for enhanced escape abilities and auditory adaptations to increase hearing acuity [31,54,55]. However, the species are known to vary in their response to risk, with the larger species being more responsive to changes in both patch quality and risk [41,56]. Similarly, the foraging patterns differ between the species, the larger applies a cream-skimming pattern, only collecting easily found grain in risky conditions, and the smaller applies a crumb-picking strategy in which they dig and search for seeds more thoroughly [42].

The gerbils were collected in the Mashabim Dunes (N 31_0’14.531’’, E 34_44’47.31’’), some 30km from the experimental enclosure. The barn owls were donated to the lab from a rescue center. For this experiment we used *Cerastes gasperetti* snakes that were trapped in the vicinity of the experimental enclosure, and housed in the zoological collection at Midreshet Ben-Gurion, Israel. All the animals were held in climate-controlled animal-rooms at the Blaustein Institutes for Desert Research within 300 m of the experimental vivarium. Both the feral dog (*Canis lupus familiaris*) and the feral cat (*Felis catus*) were trapped (adopted) in the Sde Boker environment and were loaned for the experiment by their owners.

### Experimental Set-Up

### Animals

The measurements were collected for each species in the absence of direct interspecific competition. This allowed us to make a comparative study of the effects of predation risk in the exact same setup in the absence of competition stressors. To best allow for comparison across gerbil species, we populated the vivarium with roughly the same biomass of rodents corrected for metabolic rate, leading to: 24 Allenby’s gerbils, 16 Greater Egyptian gerbils. In each of the experiments we used a 1:1 mix of adult males and females to avoid sex-based, and age-based biases and a stage structured predator prey dynamic [57]. We tracked depredation using subcutaneous RFID PIT tags which were identified by scanning the owl pellets with a reader. All other predators were muzzled to avoid physical harm to the gerbil. Depredated individuals were replaced on the following night.

### Dates

Each experiment was conducted for 100 days starting April 20^th^ 2004 with GA being on one side of the vivarium and GP on the other.

### The Enclosure

We conducted our experiments in a semi-natural, outdoor enclosure (vivarium 17 × 34 m) located on the Sede Boker Campus of Ben-Gurion University, Blaustein Institutes for Desert Research, Midreshet Ben-Gurion, Israel (N 30 51’ 25.978’’, E 34 46’ 51.284’’). The vivarium was divided into two halves. It is covered with a wire net 4.5 m high such that aerial predators can be flown within. Our vivarium mimics a dune habitat with a layer of 10 cm of sand covering a loess-clay subflooring with escape buried to 1 m below ground. To replicate the sparse vegetation cover in semi-stabilized desert dunes we created heterogeneity by adding 16 trellises in a randomized array with at least 2 m separation from the next trellis. Each trellis (80 × 50 × 15 cm) at each station was topped with a pile of cut brush to create a spherical ‘‘bush’’ approximately 1m in diameter. The environment beneath each trellis provides a sheltered environment mimicking a bush microhabitat, while the space between trellises replicates the open space occurring between vegetation clumps in the gerbils’ natural environment.

### Data Collection

We measured gerbil foraging using a randomized grid of 32 patches sieved nightly to obtain a GUD measurement. Each patch (38 × 28 × 8 cm) held 3 liters of sand and was stocked each evening with 3 g of millet. At sunrise, each patch was sieved and remaining seeds weighed to 1/100 g and logged for analysis. Daily, patches that were set up as covered patches (bush) were set outside the trellis for the following collection night as open, and vice versa. The data sampling in this experiment combined two related experiments, which used the same predators. This manuscript is a reflection of a meaningful subset of data collected. However, this resulted in an uneven number of repetitions for each predator treatment we used.

For each of the gerbil species, in accordance with each of the predator treatments and combinations of predators, two nights were sampled for each moonphase. This resulted for 8 nights of sampling for every predator alone and 4 nights for every predator combination. Additionally, 8 control nights were sampled for each rodent species in accordance with the same moonphase regimen. Because of the experimental set-up, randomization, and removal of additional data we deemed more effective to analyze separately, three of the predator combinations (cat-owl, cat-snake, and owl-snake) were run for an additional 2 nights. We randomized the predator sequence so that no two treatments would follow each other on two consequent moonphases. This means that the control nights were interspersed between other treatment nights. On nights when a single predator was used, the predator would be introduced to the vivarium once every 60 minutes from sunset to sunrise. For the snake treatment, sand imbued with the natural musk of the snakes from their enclosures was spread around the foraging stations on the same schedule. Based on the lunar calendar we marked the moonphase associated with the night of collection. Of the 64 patches, 32 were set as “bottom” trays and 32 as “full” trays. For bottom trays, we mixed the seeds into half of the sand, spread that sand over the tray, and then placed the remaining sand without seeds on top. This created a more complex tray in which the seeds were in higher concentrations, but harder to access. For full trays, we simply mixed all the seeds into all of the sand and spread that across the tray. Such setups allow us to gauge apprehension by comparing changes in selectivity [58,59]. Moonlight ambient illumination and the temperature were recorded using digital meters.

### Data Availability

The data used for this experiment available at Mendeley Data Repository [60]

### Data analyses

We separated the cumulative data collected in the experiment into two data sets: single predator data (including the control) and multi-predator data (using only nights with predator encounters every 60 minutes). We then ran analyses on the multi-species data set both as a single unit, and separately for each of the gerbil species. Figures were generated using the open source Jamovi project [61], running on R, while the statistical analyses were conducted in Systat13.

### Single predator

We ran a series of t-tests for both gerbil species combined, and for each apart. These tests compared whether the gerbils responded to the risk associated with each predator species as unique and different from the same environment in absence of predators.

### Multi-predator

To address differences between predators, we ran a General Linear Model (GLM) analysis for the data collected from the experiment using both gerbil species. We increased the explanatory value of the models, by adding interactions shown significant in a multi-factorial ANOVA that had very low explanatory power, and made sure to include all biologically meaningful interactions. The factors we used were gerbil species (GS), predator treatment (PT-type of predator present on the night of), moonphase, resource distribution (RD; clumped or dispersed resources – at the bottom of a tray or mixed throughout), and microhabitat (open and bush). For all variables and interactions found statistically significant in the model (p-values below 0.05) we ran post-hoc Tukey Tests of Honest Significance Difference (HSD).

To ease in the interpretation of post-hoc analyses, we re-ran multi-factorial ANOVAs for each of the gerbil species separately. Both these models used predator treatment, moonphase, Resource distribution and microhabitat as independent variables and added covariates of ambient temperature and moonlight intensity. Post-hoc Tukey HSD pairwise comparisons were calculated for all significant factors and interactions.

### Interpretation of data

To measure the cumulative responses of various predators, we must interpret the effects of the predators on each other, through the behavior of their prey as our experiments minimize the actual depredation our predators were allowed to perform (through restricting the predators). We use quitting harvest rates of the gerbils as gauged by giving-up densities (GUDs) in artificial patches to measure perceived risk [62,63]. We attributed patches with greater foraging activity (lower GUDs) as areas perceived as safer, and patches with higher GUDs as areas perceived as risky in accordance with previous publications[63]. The comparison of the mean GUDs for a single predator with the combination including another predator will allow us to assess the impacts of the predators on each other. By calculating a change in foraging dynamics dGUD (GUD of single – GUD of combined predators) we can expect a few patterns (Table 2). Because of predator microhabitat preference, our prediction is that facilitation will occur between the snake treatment and any of the other predators. This will result in a negative dGUD as the overall perceived risk in the environment will be higher when the two predators are present. Additionally, because we used musk in lieu of free-ranging snakes, it is likely that the response to the risk from active predators will swamp the risk from the snakes entirely. Last, if the predators facilitate each other, then the overall dGUD should be zero or positive, as the foraging in the entire experimental arena should not decline as it simply equalizes between the microhabitats [33].

**Table 2.** Inference table associating gerbil foraging with likely predator relationship. Replacement is only likely with low lethality predators, or in this experiment where an olfactory-cue was used in lieu of using live snakes.

| Main Predator | Predator Combination | Predicted Likely Outcomes |  |  |
| --- | --- | --- | --- | --- |
|  |  | dGUD if Interference | dGUD if Facilitation/<br>Cumulative Effects | dGUD if Replacement<br>(Likely focus predator) |
| Snake | Cat |  | = between microhabitats/- ↓ | -↓ (Cat) |
|  | Dog |  | = between microhabitats /- ↓ | -↓ (Dog) |
|  | Owl |  | = between microhabitats /- ↓ | -↓ (Owl) |
| Owl | Snake |  | = between microhabitats /- ↓ | +↑ (Owl) |
|  | Dog | + ↑ |  |  |
|  | Cat | + ↑ |  |  |
| Cat | Dog | + ↑ |  |  |
|  | Owl | + ↑ |  |  |
|  | Snake |  | = between microhabitats /- ↓ | =/+↑ (Cat) |
| Dog | Cat | + ↑ |  |  |
|  | Owl | + ↑ |  |  |
|  | Snake |  | = between microhabitats /- ↓ | =/+↑ (Dog) |

### Predator housing and control

Efforts were made to reduce predator strikes on gerbils: owls were flown tethered to a rope and were fed prior to the experiment, the muzzled dog ran free in the enclosure, the cat was on a harness and tether but allowed to roam freely, the tether was used to prevent the cat from making fatal strikes. As snakes were hard to control, sand imbued with their musk from their terraria was used as an olfactory cue instead.

## Results

In a simple t-test, all predators were perceived as giving rise to greater risks than the control, with exception to risk posed by owls. Allenby’s gerbils (GA) categorically associated the risk from all treatments as greater than the control, whereas the Greater Egyptian Sand Gerbils (GP) did not differentiate the snake treatment from the control and associated the owl treatment as lower risk than the control (Fig 1, Table 3). None of the treatments were perceived as a very high risk, as even the highest giving-up densities (means) were at 1.71g±0.03g, meaning that more than one third of resources were consumed in patches on a nightly average. While low, the variation seen here does not stray from published observations, and is in line with natural occurring intra-species variation.

**Fig 1.**
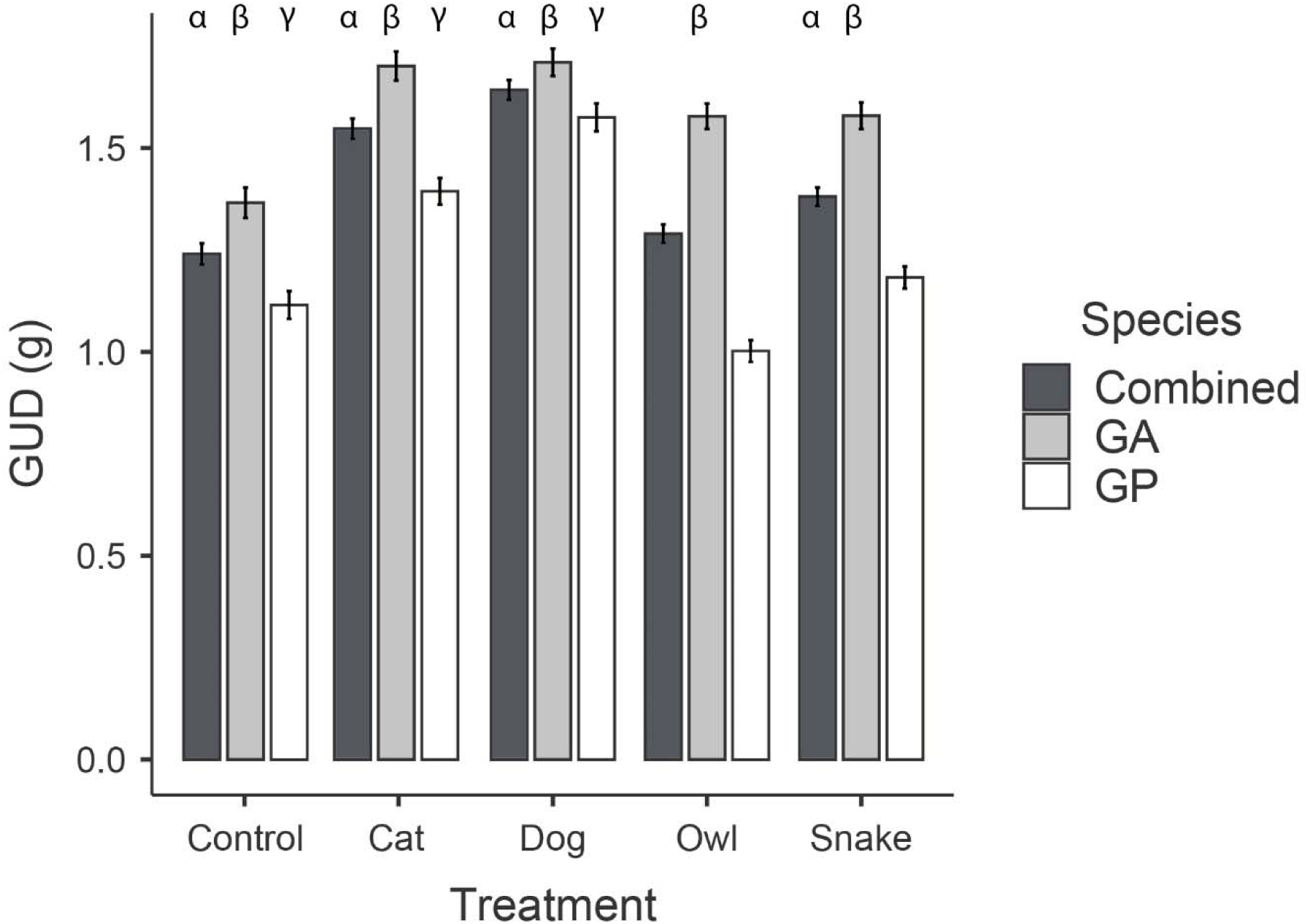
Foraging activity of each of the gerbil species and both combined in isolated, single predator treatments. α dictates a significant pairwise comparison (between control and each predator) for the combined gerbil species, β for the smaller gerbil (GA), and γ for the larger species (GP) corresponding to Table 3. The patterns shown here suggest a greater selectivity and “evaluation” of risk performed by GP than GA.

**Table 3.** T-test comparisons between the control and every single predator treatment, for the combined data set, and each of the species separately.

|  |  | **Both Gerbils** |  |  | **GA** |  |  | **GP** |  |  |
| --- | --- | --- | --- | --- | --- | --- | --- | --- | --- | --- |
| Group 1 | Group 2 | t-critical | df | P-value | t-critical | df | P-value | t-critical | df | P-value |
| Control | Snake | -4.03 | 1342 | <0.001 | -4.21 | 670 | <0.001 | -1.55 | 670 | 0.122 |
| Control | Owl | -1.44 | 1342 | 0.162 | -4.32 | 670 | <0.001 | +2.62 | 670 | 0.009 |
| Control | Cat | -8.33 | 1278 | <0.001 | -6.33 | 638 | <0.001 | -5.74 | 638 | <0.001 |
| Control | Dog | -11.11 | 1278 | <0.001 | -6.74 | 638 | <0.001 | -9.16 | 638 | <0.001 |

The GLM comparing the data collected for both gerbil species resulted in three significant single-factors and seven two-way interactions (Table 5). The two more multi-factorial ANOVAs for GA and GP separately found predator treatment, moonphase and microhabitat, and both covariates significant as well as the interactions of PT X moonphase and RD and moonphase. The three-way interaction yielded significant but hard to interpret results (SI 1,2). GA decreased foraging with increased temperature, while GP increased foraging as temperatures rose. No clear discernible patterns emerge for the moon illumination (SI 3). Regression analyses were non-significant and relatively low correlations were observed with Pearson Correlation Indices for moonlight illumination were -0.1 and 0.0 for GP and GA respectively. For temperature the correlation indices were 0.034 and 0.08 for GP and GA respectively.

GP foraged to lower GUDs than GA, with means being 1.694±0.02 and 1.489±0.02 g respectively. Both gerbils perceived the bush microhabitat as safter (lower GUDs). The means were 1.465±0.016 and 1.718±0.017 g for bush and open microhabitats respectively. Gerbil response to moonphase was not uniform for both species. GA foraged less on the full and waxing periods of the month, and had lower GUDs on the waning and new moons. The mean GUDs were 1.709±0.033, 1.751±0.030, 1.735±0.033, 1.521±0.033 g for the new, waxing, full, and waning moonphases respectively. GP foraged the least on the new and waning moonphases with GUDs being 1.521±0.033, 1.485±0.030, 1.418±0.033, 1.553±0.033 g for the new, waxing, full, and waning moonphases respectively.

Predator type (PT) yielded significant patterns in the two gerbils combined dataset, and a less impactful relationship for each separately (Fig 2). Tukey HSD tests revealed that GA response to predator treatments was a lot less definitive than with GP (Figs 2B, and 2C respectively, SI 4,5). Neither the RD, nor the interaction of moonphase with PT yielded few clear patterns of biological significance.

**Fig 2.**
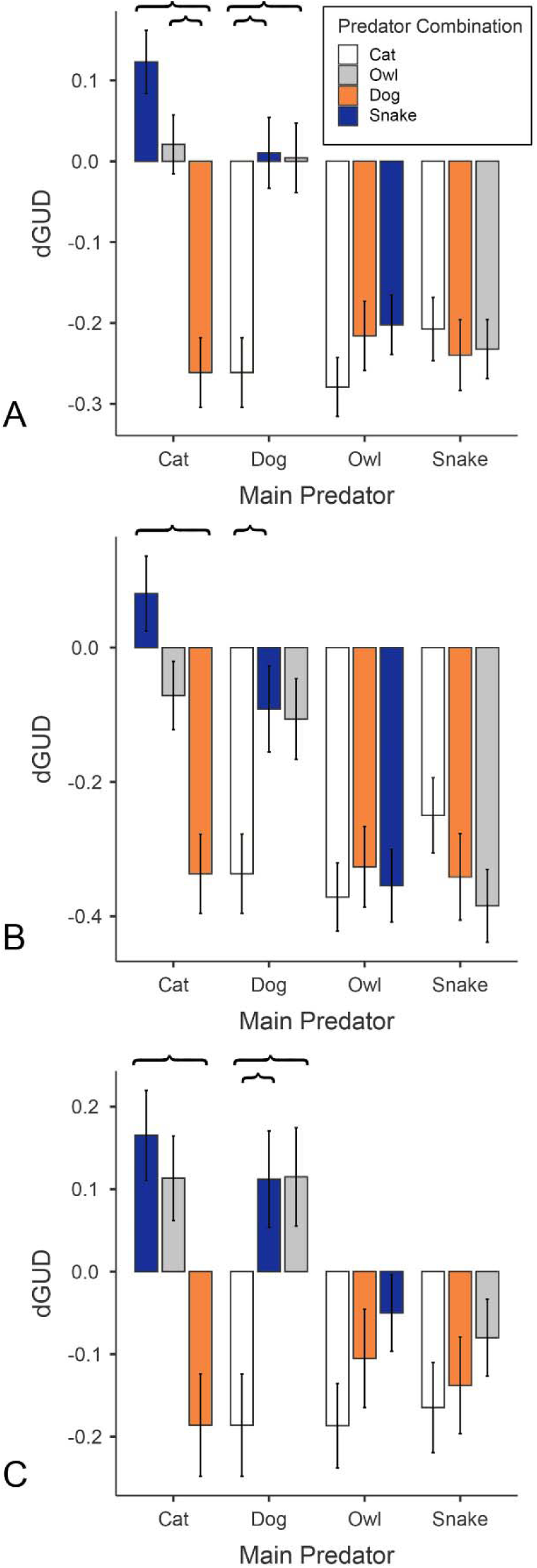
Gerbil foraging adjustment in response to perceived risk from combinations of predators. The bars represent the mean change in foraging from a treatment with a single predator and a combination of that same predator with others. (A) combined species dataset, (B) GA, and (C) GP. If the adjustment is positive (*i.e.,* combined predator GUD was lower than the single predator) the trend is suggestive of predator interference. When the adjustment is negative, the gerbils foraged less with the combination of predators than the single predator, thus suggesting a pattern of facilitation or accumulation of risk. Bold bars delineate post hoc significant pairwise comparisons (Tukey HSD) between the single predator and the combination with another. The brackets delineate significant difference between predator combinations. Asterisks represent one tailed significance (P-values between 0.05-0.1). The combined data suggests opposite patterns than observed for each species independently.

Overall, the impact of the snake treatment was negligible for both species suggesting that the gerbils were likely ignoring the olfactory cues in presence of all live predators except for the owl (Fig 3). For GA, the exposure to the dog resulted in a cumulative trend that was not statistically significant. The owl treatment was always replaced by the risk of the dog, leading to higher GUDs when the dog was present. The risk from the dog and cat treatments overall caused the largest decrease in foraging from all the predators, but were comparable to each other. The dog treatments resulted in mean GUDs of 1.61±0.037, 1.681±0.052, 1.539±0.052 g for both gerbils, GA and GP respectively. The cat, resulted in GUDs of 1.685±0.037, 1.720±0.052 and 1.650±0.052 g respectively. To better understand the predator interactions however, we examined the interaction of predator treatment and microhabitat, but only for both species combined, as the three-way interaction did not prove to be significant (Table 4, Fig 3).

**Fig 3.**
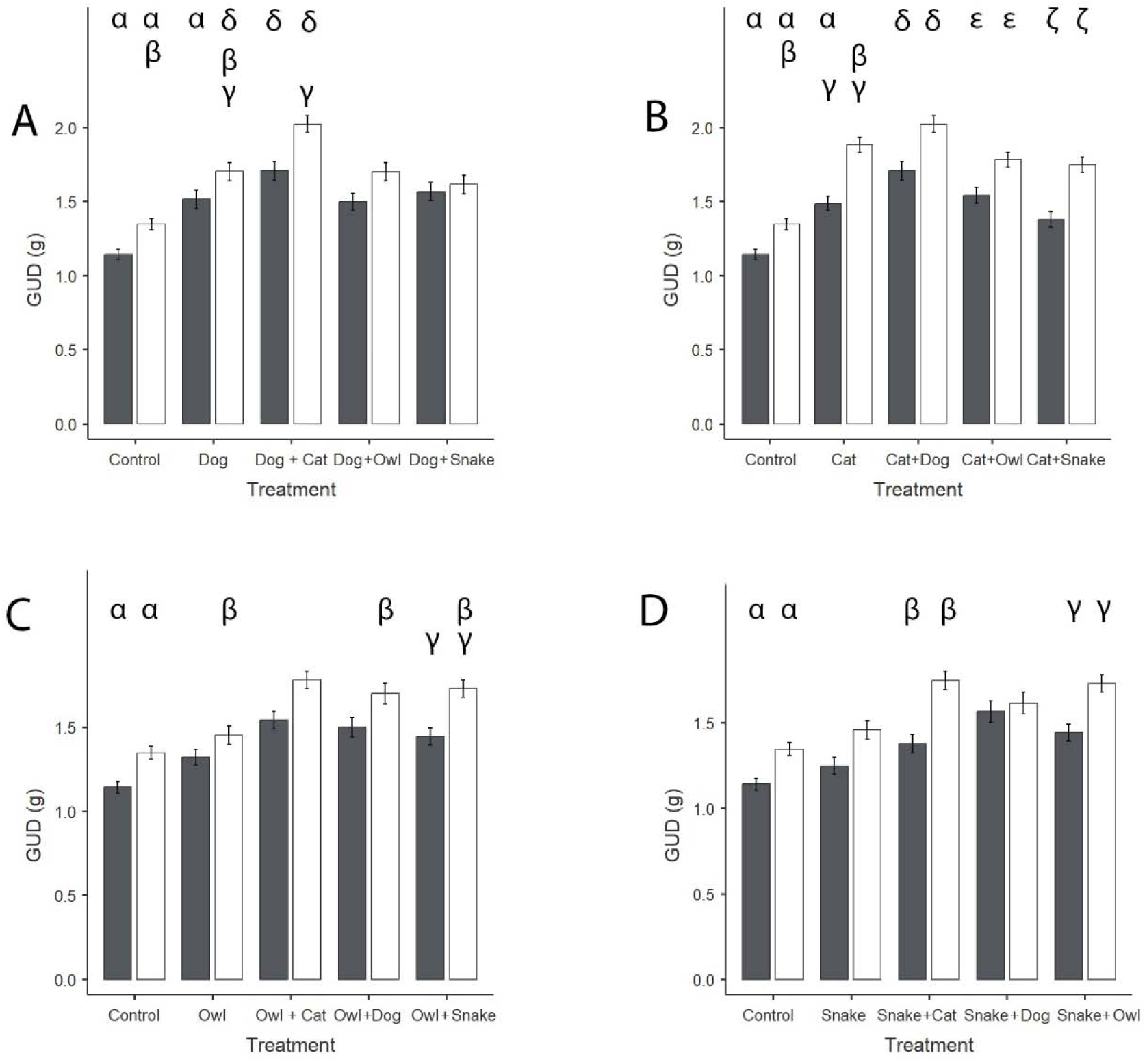
Combined Gerbil foraging (GUD±SE) as influenced by predator type and microhabitat (bush/cover in gray and open in white). Each panel focuses on a different predator and the combination with other predators where (A) focuses on the Dog, (B) the Cat, (C) the Owl, and (D) Snake musk. Greek letters represent significant Tukey Pairwise Comparisons, read from left to right for each panel.

**Table 4:** ANOVA table for GLM using data from both species (N= 2752, multiple R= 0.51, and multiple R^2^ = 0.26)

| Source | Type III<br>SS | df | Mean<br>Squares | F-Ratio | p-Value |
| --- | --- | --- | --- | --- | --- |
| Gerbil Species (GS) | 27.483 | 1 | 27.483 | 79.288 | 0.000 |
| Predator Treatment (PT) | 50.070 | 9 | 5.563 | 16.050 | 0.000 |
| Moonphase | 1.139 | 3 | 0.380 | 1.095 | 0.350 |
| Resource Distribution (Clumped/ Evenly<br>Mixed) (RD) | 0.031 | 1 | 0.031 | 0.090 | 0.764 |
| Microhabitat | 41.602 | 1 | 41.602 | 120.021 | 0.000 |
| GS X PT | 9.019 | 9 | 1.002 | 2.891 | 0.002 |
| Moonphase X GS | 6.922 | 3 | 2.307 | 6.656 | 0.000 |
| Microhabitat X GS | 22.670 | 1 | 22.670 | 65.401 | 0.000 |
| Moonphase X PT | 72.831 | 27 | 2.697 | 7.782 | 0.000 |
| Microhabitat X PT | 6.701 | 9 | 0.745 | 2.148 | 0.023 |
| Microhabitat X RD | 3.834 | 1 | 3.834 | 11.060 | 0.001 |
| RD X GS | 0.256 | 1 | 0.256 | 0.739 | 0.390 |
| GS X PT X Moonphase | 59.957 | 27 | 2.221 | 6.407 | 0.000 |
| PT X Moonphase X RD | 18.991 | 27 | 0.703 | 2.029 | 0.001 |
| Error | 911.965 | 2,631 | 0.347 |  |  |

**Table 5.** Three-way interactions post hoc Tukey Tests of Honestly Significant Difference for A. Gerbillus andersoni allebyi, and B. Gerbillus pyramidum. Only significant interactions are listed, and * delineates one tailed distribution significance

| GA |  |  |  |  |  |  |  |
| --- | --- | --- | --- | --- | --- | --- | --- |
| # | Fixed Variable | Category | Dynamic Variable | Relationship |  |  | P-value |
| 1 | Moonphase | Full | Predators | Dog | > | Snake | 0.022 |
| 2 | Moonphase | New | Predators | Dog+ Snake | > | Snake | <0.001 |
| 3 | Moonphase | New | Predators | Owl | > | Snake | <0.001 |
| 4 | Moonphase | Waning | Predators | Cat + Dog | > | Cat + Snake | <0.001 |
| 5 | Moonphase | Waning | Predators | Dog + Owl | > | Owl | 0.009 |
| 6 | Moonphase | Waxing | Predators | Dog + Cat | > | Dog + Snake | 0.009 |
| 7 | Moonphase | Waxing | Predators | Dog + Cat | > | Dog | <0.001 |
| 8 | Moonphase | Waxing | Predators | Dog + Cat | > | Dog + Snake | 0.001 |
| 9 | Predators | Cat + Snake | Moonphase | Full | > | Waning | <0.001 |
| 10 | Predators | Dog | Moonphase | New | > | Full | 0.023 |
| 11 | Predators | Dog | Moonphase | Full | > | Waxing | <0.001 |
| 12 | Predators | Dog+ Snake | Moonphase | New | > | Waxing | 0.013 |
| 13 | Predators | Owl | Moonphase | Waxing | > | New | 0.059* |
| 14 | Predators | Cat + Owl | Moonphase | Waxing | > | Waning | 0.031 |
| GP |  |  |  |  |  |  |  |
| # | Fixed Variable | Category | Dynamic Variable | Relationship |  |  | P-value |
| 1 | Moonphase | Full | Predators | Cat | > | Dog | 0.058* |
| 2 | Moonphase | Full | Predators | Cat + Owl | > | Dog | 0.044 |
| 3 | Moonphase | Full | Predators | Cat + Dog | > | Cat + Snake | <0.001 |
| 4 | Moonphase | Full | Predators | Cat + Dog | > | Dog | <0.001 |
| 5 | Moonphase | Full | Predators | Owl | > | Dog | <0.001 |
| 6 | Moonphase | New | Predators | Cat | > | Owl | <0.001 |
| 7 | Moonphase | New | Predators | Cat + Owl | > | Owl | <0.001 |
| 8 | Moonphase | New | Predators | Dog | > | Dog + Snake | 0.048 |
| 9 | Moonphase | New | Predators | Dog | > | Owl | <0.001 |
| 10 | Moonphase | New | Predators | Dog + Cat | > | Dog + Snake | <0.001 |
| 11 | Moonphase | New | Predators | Dog + Owl | > | Owl | <0.001 |
| 12 | Moonphase | Waning | Predators | Cat | > | Snake | <0.001 |
| 13 | Moonphase | Waning | Predators | Cat + Owl | > | Dog + Owl | 0.008 |
| 14 | Moonphase | Waning | Predators | Cat + Snake | > | Snake | <0.001 |
| 15 | Moonphase | Waxing | Predators | Dog + Owl | > | Dog + Cat | 0.022 |
| 16 | Moonphase | Waxing | Predators | Dog | > | Owl | <0.001 |
| 17 | Moonphase | Waxing | Predators | Dog | > | Snake | <0.001 |
| 18 | Predators | Cat + Snake | Moonphase | Waning | > | Full | <0.001 |
| 19 | Predators | Dog | Moonphase | New | > | Full | <0.001 |
| 20 | Predators | Dog | Moonphase | Waxing | > | Full | <0.001 |
| 21 | Predators | Dog+ Snake | Moonphase | Waxing | > | New | 0.046 |

The two ground foraging predators seemed to have elicited the largest “fear” response in the gerbils, corresponding with highest mean GUDs. Between the two predators, the cat was perceived as a much greater threat than the dog (Fig 3 A&B). The change in foraging in the open microhabitat decreased by 10.8% of the total available food from the dog treatment to the dog and cat combined. The decrease was only 4.6% from the cat to the cat + dog combination. Neither the snake treatment alone, nor owl treatment were substantially different from the control in this experiment, in neither the bush nor the open microhabitat (Fig 3 C&D). For the owl, the open microhabitat became significantly riskier only in combination with the dog and the snake, but not with the cat (an anomalous result). Last, the snake treatment resulted in very little differences from the control. However, when combined with the dog, both microhabitats were equally risky pointing to facilitation between the two predators.

## Discussion

This experiment highlights and strengthens a few previous finds, namely that not all predators exude the same intensity of predation risk on foragers [29,64,65], and that different forager species-based on their unique adaptive strategies-do not respond equally to the same risks [66,67]. Meanwhile, our data also supports observation made by numerous authors that the suggest that intensity of a predator cue affects the response by the prey [68], and that the presence of various predators simultaneously have cumulative impacts and not substitutive effects. However, the perceived risk that a predator exudes on the prey will dictate whether the effect of the additional risk factor is negligible or cumulative.

### Foraging strategy matters

Stronger interference competitors that employ a cream-skimming foraging strategy appear to be more attuned to the specifics of risk in their environment. GPs were much more specific in their management of light (moonphase) and predator activity than the GA were. This suggests that when foragers have the ability to exclude their competition, they will use resources in a more discerning manner than a lesser competitor [39]. A number of hypotheses have been generated over time that explain why the larger competitor tends to “give-up” sooner than the smaller one. The leading ones include that the larger prey have greater value to the predator and thus draw greater focus from the predator [69]; another is that larger foragers have higher energetic foraging costs due to the larger metabolic costs associated with greater body mass [20]. Regardless of explanation, the evidence collected herein supports this observation. The consequence of the sensitivity of the larger competitor is that it “opens up” time, space, and resources for the lesser competitor who is less discerning [70,71].

The observations relating to the smaller competitor, GA, suggest a different pattern of behavior than its larger counterpart. In congruence with former observations, the GA accept what they must as the weaker competitor. They take more risks [20], utilize suboptimal times [67] and suboptimal substrates [53,71]. Last, they forage in patches that have much lower energetic value (Kotler et al. 1993b, Ben-Natan et al. 2004). The two major observations supporting this dynamic in our data were a hyper focus on the greatest risk predators in the experiment, where they foraged more, and took risks with all predators except the highest risk (dog or cat). Additionally, the support of the suboptimal times, the GA preferred the “in-between” times when the moon was waning and waxing especially at times that the risk was greatest [72].

### Predator treatments

The patterns of facilitation, interference and cumulative risks derived from the combinations of predators significantly deviated from our expectations. The response to olfactory cues were smaller than the responses to live predators, and even a live predator did not always differ significantly from the control treatment (as seen in the example of the owl). Below we aim to break down the possible drivers of the interactions we observed.

### Predation risk administration

It is not surprising that a predator cue in absence of the actual predator, was perceived as a lower risk than live predators. Many studies have tested the use of olfactory cues of predators as deterrents to control agricultural damage and found conflicting outcomes. This scenario, where the gerbils underestimate the risk from the snake musk, also aligns with a significant body of literature that suggests that olfactory cues result in less impactful results from prey. A couple of examples include Tasmanian wallabies having little response to dingo urine [73] and squirrels in Illinois not responding to red fox urine [74,75]. Moreover, the debate about using olfactory cues to simulate predation has been ongoing for decades. Banks et al. [76,77] have shown that the response to olfactory cues can vary significantly, especially when the source of the cue is not fresh. They also argue that these types of cues are used in a subjective manner by the foragers, for example bush rats in their experiments had less response to old urine when exposed to fresh urine on the same night. Olfactory cues have also been shown to be interpreted by prey differently based on ambient temperature and air humidity [78]. Lastly, foragers have even been found to use olfactory cues, of both predators and competitors, as camouflage from live predators[68].

We mention all these challenges to raise the caveat that we interpret our results to suggest that the gerbil responses to the olfactory cues for the presence of snakes, based on musk-imbued sand, do not support the view that gerbils would interpret the scent as a sign of predation risk. However, the gerbils did show an interaction between these cues and the presence of other predators, namely owl and dog. In addition, the responses were much greater for the GPs than for the GA, evidence supporting the above claims on variation between the species and their competitive foraging strategies.

Hunting strategy matters and impacts prey’s risk perception.

Foraging has been shown to decrease based on various predator related characteristics, namely the lethality of the predator, but potentially more importantly, based on perceived risk which at times may veer from the actual lethality of the predator [79,80]. Thus, as a generalization, prey often overestimate the risk posed by a predator, as the cost of under-estimating the risk is significantly greater than the opposite [81–84]. Additionally, the information a prey species has of their predator’s behavior, innate or learned, will change the behavioral tactics they will employ to avoid that risk. While the most commonly reported is a shift in microhabitat use, the entire spatial and temporal distribution may change based on the type of predator the foragers interact with. For example, desert rodents from multiple continents have been shown to reassess risk distribution, sometimes referred to as a landscape of fear, and refocused their anti-predator strategies to the predator they perceived as more lethal. In that example the greater risk was provided by an owl that overshadowed live snakes [65].

Our observations in this experiment suggest a general agreement between the two foragers. Both the gerbils foraged the least in the presence of the active ground hunting predators, followed by the owl and the snake musk having a significantly lower impact on their behavior. Similarly, the cat was perceived as a greater risk than the dog as evident by the significant decrease in foraging when the combination of the cat and dog are compared to the isolated dog treatment. However, when comparing the combined predators with an isolated cat, the pattern is cumulative but not significant.

Predator facilitation, interference, and cumulative effects.

The interactions between two predator treatments overwhelmingly showed cumulative effects. Where the interaction was not significant, this can be interpreted as additive interactions. Where the interaction was less than additive (significant positive dGUD) we can interpret the interaction as interference. When the interaction was more than additive (significant negative dGUD) we interpret the result to be facilitation. The gerbils of both species foraged less in combinations of predators over the control and over each predator alone. However, the specific interactions do shed light on a couple of patterns. Where interference between predators was observed, as measured through increased foraging in the combined treatment, it was never statistically significant, and thus the interpretation results in a conclusion that these are all additive interactions. Additive patterns were more likely in the GP population than in GA and were more likely in predators of lower lethality (owl and snake) and especially when these predators hunt using different strategies as in the snake and the cat (present in both species). A plausible explanation for why these appear to exist is that different parts of the enclosure are associated with the risk of specific predators. For example, both Merriam’s kangaroo rats (*Dipodomys merriami*) and GAs have been shown to identify areas of risk of predators they prioritize [30] and change their geographic distribution within experimental enclosures in response. The kangaroo rats had identified the locations where snakes were present and depleted the available forage in other locations, meanwhile GAs avoided a transact of land corresponding to the flight path between two perches favored by the barn owls but were shown to be willing to risk the physical space where snakes were observed. Having the combination of multiple predators here, drives the gerbils to search for resources in environment they did not previously forage, and thus increasing their harvest intake to compensate for increased movement activity [48].

The interplay of predator hunting strategies was expected to yield more significant habitat selections in the gerbils than we observed. The clearest of textbook examples for predator facilitation is that of an active-searching predator, like an owl, pushing the foragers towards ambush predators awaiting them under bushes. The presence of ambush predators alternatively encourages the foragers to avoid the refuge for the exposed environment where the owls can pick them off [33]. As mentioned above, our ambush predator treatment, in the form of snake musk, was not as effective as we had planned and resulted in the bush habitat being safer in every interaction where an owl was present. The dog treatment overall interplayed similarly to that owl, as a larger mesocarnivores that would predominantly impose risk in the open spaces and could not fit under the trellises. The cat on the other hand equalized the risk in both microhabitats, especially when interacting with the other active searching predators. This may also explain the overall greater impact that are perceived by feral cats, and their known impact to small vertebrate biodiversity in arid zones [85].

Despite our expectation, the theory of predator facilitation did hold up when we look at the microhabitat use in this experiment. This perhaps is a result of the lack of significance of the three-way interaction of gerbil species, predator treatment and microhabitat. Therefore, we were only observing interactions of the combined data set for both species. Here, the only example of facilitation was not between the cat and snake as we had initially expected, but between the dog and the snake. In that treatment combination the gerbils equalized the resource use between the microhabitats. To our best interpretation, this comes from the general behavioral pattern both gerbil species were observed to follow where they shift their focus to the greatest risk in the environment [30]. While all the other interactions appear to be facilitative and cumulative with the foraging decreasing in the open microhabitat only, it would be very difficult to tease out which of the two each interaction represents. However, the same pattern of shift of attention leads us to assume that most of the interactions are cumulative.

### The anthropogenic human-commensal predator, conservation implications

Our use of feral, human commensal species of predators allows us to comment on an unexpected aspect of conservation importance. Significant numbers of papers focus on the impact that feral cats specifically [85–88] and human commensal predators in general [89–91] have on biological communities globally. Our observations were both encouraging and support the overall fear of the exploding populations of feral cats, a politically explosive topic in arid zones, such as Australia [92], the Middle East and North Africa [93] and on arid islands [94]. The encouragement we observe comes from the fact that the gerbils clearly understood the cat to be the greatest threat to them. In addition, the gerbils interpreted the risk from the cats and dogs correctly and responded by shifting their microhabitat preference. Thus the overall impact these feral predators may have on their populations is curbed in comparison to the impacts they have on naïve prey on islands and the Australian continent [95,96].

In addition, the impact the dog had on the foraging of the gerbils surprised us. Predominantly, aerial predators, and specifically barn owls, tend to garner the most anti-predator focus in gerbils [35]. It is unusual to find ground dwelling, active searching predator, as the main risk factor for desert granivores. The fact that the dog, despite being perceived as a lower risk than the cat, still resulted in greater apprehension by the gerbils, suggests that human commensal predators have a significant potential to impact community-wide interactions around the globe.

Community-wide impacts of feral cats and dogs have predominantly focused on consumptive and injury-based effects, with strong emphasis on the naiveté of prey [49,51,94,95,97]. Studies on feral cat impacts far outnumber those on feral dogs[98]. The study of community-wide impacts of feral dogs has been mostly limited to the role of dingoes replacing apex predators in Australian arid zones. Two major factors differentiate this case from the present studies: first, dingoes act as apex predators in Australia, while in our study, feral dogs functioned as meso-carnivores within Negev and Saharan Desert ecosystems; second, management efforts for dingoes—and therefore research on them—far exceed those for feral dogs in other locations. Meanwhile, reviews of prey responses to dingoes, albeit focused on larger prey species, indicate that responses vary based on cue type and recent exposure to the predator. In other feral dog populations, most research has focused solely on consumptive and injury-based impacts.

However, recent evidence challenges the assumption that native prey are naïve to introduced predators[98,99].While previously attributed biodiversity demise in arid zones was attributed to naïveté of the prey, more recent studies show that small mammals in arid regions are rapidly adapting to recognize the presence of feral and invasive predators. The exception lies in feral cats [98]. Our observations here stand at odds with these observations from the Australian studies. The gerbils clearly acknowledged the risk posed by the cats in their behavioral choices. We offer two interpretive reasons for this difference. First domestic cats are believed to have been domesticated from an ancestral species *Felis libyaca* in this region of the world[100], suggesting a much longer period of habituation with both the feral dogs and cats. Second, we expect a certain evolved recognition given the diversity of native felid and canine mesocarnivores in the habitat where the gerbils were collected, namely jackals, sand foxes, red foxes, caracals and sand cats.

While not groundbreaking, our observations have significant implications for conservation and wildlife management. Predominantly, we can acknowledge the need for strategies beyond predator removal, which is economically and ecology costly and perceived as a controversial practice by the general public [101]. However, this raises the need to manage wildlife by enhancing natural antipredator behaviors, potentially through controlled exposure to predator cues, which should reduce the long-term impacts of the feral predators. Additionally, habitat modifications that increase cover and refuge availability can mitigate predation risk, and reduce consumptive impacts. This would support small mammal populations in arid environments where exposure to predators increases in movement between shelters [15]. By integrating these approaches with traditional predator control measures, conservation efforts can better safeguard biodiversity in landscapes where human-commensal predators exert disproportionate influence.

## Conclusion

The experiment was designed to tease out the interactions of various predator types on the foraging behavior of small, wild, desert mammals. Through the behavior of the gerbils, the facilitation and interference between the predators was tested. The interactions between predators proved to be predominantly cumulative. Facilitation was found to be indisputable between lower-lethality snake olfactory cues and the dog. The impacts of ground foraging, human-commensal species, was shown to draw the attention of the gerbils over predators that naturally occur in their environment, such as a barn owl and a horned viper. This also supports a tendency of habituation to these predators as integral parts of the predator community, and not a novelty risk to which prey species are naïve. The tendency of the gerbils to focus their apprehension on the greatest risk in the environment supports the use of single predator type titrations favored in behavioral common-garden experiments, but potentially only true to these specific species. Therefore, these findings might or might not generalize to other arid-zone small mammals or other introduced carnivores.

## Acknowledgements

The research was funded by a United States–Israel Binational Science Foundation grant #1999– 109 to BPK and JSB. We thank the Midreshet Ben Gurion Zoological Collection Reptile Collection for providing snake musk imbued bedding used in these experiments. Graphical abstract illustrated by Reza Dalvand.

## Declaration of generative AI and AI-assisted technologies in the writing process

During the preparation of this work the authors used Chat GPT in order to generate base illustrations based on personal photographs for the graphical abstract. No AI was used in writing or editing this work.

## Author Contributions

Conceptualization: Justin St Juliana and Burt Kotler, Methodology: Justin St. Juliana, Joel Brown and Burt Kotler; Formal analysis and investigation: Sonny Bleicher, Justin St Juliana, Shomen Mukherjee and Vijayan Sundararaj; Writing – original draft preparation: Sonny Bleicher; Writing – review and editing: Justin St Juliana, Burt Kotler and Sonny Bleicher, Funding acquisition: Burt Kotler and Joel Brown; Resources: Burt Kotler and Joel Brown; Supervision: Burt Kotler

## Ethical Statement

### Research involving Human Participants and/or Animals

The research was conducted in compliance with the provisions of the Israel Nature and Parks Authority, in lieu of an ethics committee established at Ben Gurion University at the time of this experiment. The animals collected in the sand dunes were collected under permit from the Israel Nature and Parks Authority Permit number 01/2003 to BPK. The procedures used in this study adhere to the tenets of the Declaration of Helsinki.

## List of Appendices

SI 1. Multi-way ANOVA for GA, including post-hoc Tukey tests for all significant variables and interactions.

SI 2. Multi-way ANOVA for GP, including post-hoc Tukey tests for all significant variables and interactions.

SI 3. Pearson Correlation matrices for the covariates against GUD for each of the gerbil species separately.

SI 4. Response of combined gerbil species to predator encounters.

SI 5. Response of gerbils to predator treatments as affected by moonphase. SI 6. Raw-data file used for analyses.

## References

1. Ale, S.B.; Whelan, C.J. Reappraisal of the Role of Big, Fierce Predators! Biodivers. Conserv. 2008, 17, 685–690, doi:10.1007/s10531-008-9324-5.

2. Laundré, J.W.; Hernandez, L.; Ripple, W.J. The Landscape of Fear: Ecological Implications of Being Afraid. Open Ecol. J. 2010, 3, 1–7, doi:10.2174/1874213001003030001.

3. Rosenzweig, M.L.; MacArthur, R.H. Graphical Representation and Stability Conditions of Predator-Prey Interactions. Am. Nat. 1963, 97, 209–223.

4. Stephan, J.G.; Stenberg, J.A.; Björkman, C. Consumptive and Non-Consumptive Effect Ratios Depend on Interaction between Plant Quality and Hunting Behaviour of Omnivorous Predators Running Head[: Plant Quality and Hunting Behaviour Affecting Prey. Ecol. Evol. 2017, 7, 2327–2339, doi:10.1002/ece3.2828.

5. Peckarsky, B.L.; Abrams, P.A.; Bolnick, D.I.; Dill, L.M.; Grabowski, J.H.; Luttbeg, B.; Orrock, J.L.; Peacor, S.D.; Preisser, E.L.; Schmitz, O.J.;, et al. Revisiting the Classics: Considering Nonconsumptive Effects in Textbook Examples of Predator Prey Interactions. Ecology 2008, 89, 2416–2425, doi:10.1890/07-1131.1.

6. Paine, R.T. Food Web Complexity and Species Diversity. Am. Nat. 1963, 100, 65–75, doi:10.2307/2678832.

7. Lima, S.L. Nonlethal Effects in the Ecology of Predator-Prey Interactions. Bioscience 1998, 48, 25–34, doi:10.2307/1313225.

8. Orrock, J.L.; Grabowski, J.H.; Pantel, J.H.; Peacor, S.D.; Peckarsky, L.; Sih, A.; Werner, E.E.; Werner, E. Consumptive and Nonconsumptive Effects of Predators on Metacommunities of Competing Prey. Ecology 2008, 89, 2426–2435.

9. Sih, A.; Bolnick, D.I.; Luttbeg, B.; Orrock, J.L.; Peacor, S.D.; Pintor, L.M.; Preisser, E.; Rehage, J.S.; Vonesh, J.R. Predator-Prey Naïveté, Antipredator Behavior, and the Ecology of Predator Invasions. Oikos 2010, 119, 610–621, doi:10.1111/j.1600-0706.2009.18039.x.

10. Schmitz, O.J.; Beckerman, A.P.; Brien, K.M.O. Behaviorally Mediated Trophic Cascades[: Effects of Predation Risk on Food Web Interactions. Ecology 1997, 78, 1388–1399.

11. Letnic, M.; Ritchie, E.G.; Dickman, C.R. Top Predators as Biodiversity Regulators: The Dingo Canis Lupus Dingo as a Case Study. Biol. Rev. 2012, 87, 390–413, doi:10.1111/j.1469-185X.2011.00203.x.

12. Beschta, R.L.; Ripple, W.J. Large Predators and Trophic Cascades in Terrestrial Ecosystems of the Western United States. Biol. Conserv. 2009, 142, 2401–2414, doi:10.1016/j.biocon.2009.06.015.

13. Kotler, B.P.; Holt, R.D. Predation and Competition: The Interaction of Two Types of Species. Oikos 1989, 54, 256–260.

14. Manlick, P.J.; Newsome, S.D. Adaptive Foraging in the Anthropocene: Can Individual Diet Specialization Compensate for Biotic Homogenization? Front. Ecol. Environ. 2021, 19, 510–518, doi:10.1002/fee.2380.

15. Bleicher, S.S.; Dickman, C.R. On the Landscape of Fear: Shelters Affect Foraging by Dunnarts (Marsupialia, Sminthopsis Spp.) in a Sandridge Desert Environment. J. Mammal. 2020, 101, 281–290, doi:10.1093/jmammal/gyz195.

16. Pavey, C.R.; Eldridge, S.R.; Heywood, M. Population Dynamics and Prey Selection of Native and Introduced Predators during a Rodent Outbreak in Arid Australia. J. Mammal. 2008, 89, 674–683, doi:10.1644/07-MAMM-A-168R.1.

17. Dickman, C.R.; Greenville, A.C.; Tamayo, B.; Wardle, G.M. Spatial Dynamics of Small Mammals in Central Australian Desert Habitats: The Role of Drought Refugia. J. Mammal. 2011, 92, 1193–1209, doi:10.1644/10-MAMM-S-329.1.

18. Abramsky, Z.; Rosenzweig, M.L.; Subach, A. The Costs of Apprehensive Foraging. Ecology 2002, 83, 1330–1340.

19. Kotler, B.P.; Blaustein, L. Titrating Food and Safety in a Heterogeneous Environment[: When Are the Risky and Safe Patches of Equal Value[? Oikos 1995, 74, 251–258.

20. Brown, J.S.; Kotler, B.P. Hazardous Duty Pay and the Foraging Cost of Predation. Ecol. Lett. 2004, 7, 999–1014, doi:10.1111/j.1461-0248.2004.00661.x.

21. Relyea, R.A. How Prey Respond to Combined Predators[: A Review and an Empirical Test. Ecology 2003, 84, 1827–1839.

22. Arditi, R.; Ginzburg, L.R. Coupling in Predator-Prey Dynamics: Ratio Dependence. J. Theor. Biol. 1989, 139, 311–326.

23. St. Juliana, J.; Kotler, B.P.; Brown, J.S.; Mukherjee, S.; Bouskila, A. The Foraging Response of Gerbils to a Gradient of Owl Numbers. Evol. Ecol. Res. 2011, 13, 869–878.

24. Mukherjee, S. Understanding the Interplay Between Time Allocation, Vigilance and State Department Foraging Behavior. (Doctoral Dissertation, Ben Gurion University of the Negev)., Ben Gurion University, 2010.

25. Berec, L. Impacts of Foraging Facilitation among Predators on Predator-Prey Dynamics. Bull. Math. Biol. 2010, 72, 94–121, doi:10.1007/s11538-009-9439-1.

26. Soluk, D.A.; Collins, N.C. Between Fish and Stoneflies[: Facilitation Interactions Synergistic and Interference among Stream Predators. Oikos 1988, 52, 94–100.

27. Embar, K.; Kotler, B.P.; Bleicher, S.S.; Brown, J.S. Pit Fights: Predators in Evolutionarily Independent Communities. J. Mammal. 2018, 99, 1183–1188, doi:10.1093/jmammal/gyy085.

28. Kotler, B.P.; Blaustein, L.; Brown, J.S. Predator Facilitation: The Combined Effect of Snakes and Owls on the Foraging Behavior of Gerbils. Ann. Zool. Fenn. 1992, 29, 199–206.

29. Bleicher, S.S.; Brown, J.S.; Embar, K.; Kotler, B.P. Novel Predator Recognition by Allenby’s Gerbil (Gerbillus Andersoni Allenbyi): Do Gerbils Learn to Respond to a Snake That Can “See” in the Dark? Isr. J. Ecol. Evol. 2016, 62, 178–185, doi:10.1080/15659801.2016.1176614.

30. Bleicher, S.S.; Kotler, B.P.; Brown, J.S. Comparing Plasticity of Response to Perceived Risk in the Textbook Example of Convergent Evolution of Desert Rodents and Their Predators; a Manipulative Study Employing the Landscape of Fear. Front. Behav. Neurosci. 2019, 13, 1–12, doi:10.3389/fnbeh.2019.00058.

31. Kotler, B.P.; Brown, J.S.; Bleicher, S.S.; Embar, K. Intercontinental-Wide Consequences of Compromise-Breaking Adaptations: The Case of Desert Rodents. Isr. J. Ecol. Evol. 2016, 62, 186–195, doi:10.1080/15659801.2015.1125832.

32. Bleicher, S.S.; Kotler, B.P.; Downs, C.J.; Brown, J.S. Playing to Their Evolutionary Strengths; Heteromyid Rodents Provide Opposite Snake Evasion Strategies in the Face of Known and Novel Snakes. J. Arid Environ. 2020, 173, 104025, doi:10.1016/j.jaridenv.2019.104025.

33. Embar, K.; Raveh, A.; Hoffmann, I.; Kotler, B.P. Predator Facilitation or Interference: A Game of Vipers and Owls. Oecologia 2014, 174, 1301–1309, doi:10.1007/s00442-013-2760-2.

34. Leo, V.; Reading, R.P.; Letnic, M. Interference Competition: Odours of an Apex Predator and Conspecifics Influence Resource Acquisition by Red Foxes. Oecologia 2015, 179, doi:10.1007/s00442-015-3423-2.

35. Kotler, B.P.; Brown, J.S.; Mukherjee, S.; Berger-Tal, O.; Bouskila, A.; Kotier, B.P.; Brown, J.S.; Mukherjee, S.; Berger-Tal, O.; Bouskila, A. Moonlight Avoidance in Gerbils Reveals a Sophisticated Interplay among Time Allocation, Vigilance and State-Dependent Foraging. Proc. R. Soc. B 2010, 277, 1469–1474, doi:10.1098/rspb.2009.2036.

36. Wasko, D.K.; Bonilla, F.; Sasa, M. Behavioral Responses to Snake Cues by Three Species of Neotropical Rodents. J. Zool. 2014, 292, doi:10.1111/jzo.12086.

37. Orizaola, G.; Dahl, E.; Laurila, A. Reversibility of Predator-Induced Plasticity and Its Effect at a Life-History Switch Point. Oikos 2012, 121, 44–52, doi:10.1111/j.1600-0706.2011.19050.x.

38. Bleicher, S.S.; Kotler, B.P.; Downs, C.J.; Brown, J.S. Intercontinental Test of Constraint[Breaking Adaptations; Testing Behavioural Plasticity in the Face of a Predator with Novel Hunting Strategies. J. Anim. Ecol. 2020, 89, 1837–1850, doi:10.1111/1365-2656.13234.

39. Abramsky, Z.; Strauss, E.; Subach, A.; Riechman, A.; Kotler, B.P. The Effect of Barn Owls (Tyto Alba) on the Activity and Microhabitat Selection of Gerbillus Allenbyi and G. Pyramidum. Oecologia 1996, 105, 313–319, doi:10.1007/BF00328733.

40. Rosenzweig, M.L. Habitat Selection as a Source of Biological Diversity. Evol. Ecol. 1987, 1, 315–330, doi:10.1007/BF02071556.

41. Ben-Natan, G.; Abramsky, Z.; Kotler, B.P.; Brown, J.S. Seeds Redistribution in Sand Dunes[: A Basis for Coexistence of Two Rodent Species. Oikos 2004, 105, 325–335, doi:10.1111/j.0030-1299.2004.12948.x.

42. Brown, J.S.; Kotler, B.P.; Mitchell, W.A. Competition between Birds and Mammals[: A Comparison of Giving-up Densities between Crested Larks and Gerbils. Evol. Ecol. 1997, 11, 757–771, doi:10.1023/A:1018442503955.

43. Kotler, B.P.; Brown, J.S.; Dall, S.R.X.; Gresser, S.; Ganey, D.; Bouskila, A. Foraging Games between Gerbils and Their Predators[: Temporal Dynamics of Resource Depletion and Apprehension in Gerbils. Evol. Ecol. Res. 2002, 4, 495–518.

44. Biewener, A.; Blickhan, R. Kangaroo Rat Locomotion: Design for Elastic Energy Storage or Acceleration? J. Exp. Biol. 1988, 140, 243–255.

45. Longland, W.S.; Price, M. V Direct Observations of Owls and Heteromyid Rodents[: Can Predation Risk Explain Microhabitat Use[? Ecology 1991, 72, 2261–2273.

46. Webster, D.B.; Strother, W.F. Middle-Ear Mor-Phology and Auditory Sensitivity of Heteromyid Rodents. Am. Zool. 1972, 12, 727.

47. Abramsky, Z.; Rosenzweig, M.L.; Subach, A. The Cost of Interspecific Competition in Two Gerbil Species. J. Anim. Ecol. 2001, 70, 561–567.

48. Makin, D.F.; Kotler, B.P. How Do Allenby ‘ s Gerbils Titrate Risk and Reward in Response to Different Predators[? Behav. Ecol. Sociobiol. 2020, 74, 1–10, 10.1007/s00265-019-2785-6 (2020).

49. Letnic, M.; Koch, F. Are Dingoes a Trophic Regulator in Arid Australia? A Comparison of Mammal Communities on Either Side of the Dingo Fence. Austral Ecol. 2010, 35, 167–175, doi:10.1111/j.1442-9993.2009.02022.x.

50. Berger, S.; Wikelski, M.; Romero, L.M.; Kalko, E.K. V; Rödl, T. Behavioral and Physiological Adjustments to New Predators in an Endemic Island Species, the Galápagos Marine Iguana. Horm. Behav. 2007, 52, 653–663, doi:10.1016/j.yhbeh.2007.08.004.

51. Carthey, A.J.R.; Blumstein, D.T. Predicting Predator Recognition in a Changing World. Trends Ecol. Evol. 2018, 33, 106–115, doi:10.1016/j.tree.2017.10.009.

52. Goodfriend, W.; Ward, D.; Subach, A. Standard Operative Temperatures of Two Desert Rodents, Gerbillus Allenbyi and Gerbillus Pyramidum: The Effects of Morphology, Microhabitat and Environmental Factors. J. Therm. Biol. 1991, 16, 157–166, doi:10.1016/0306-4565(91)90038-4.

53. Kotler, B.P.; Brown, J.S.; Oldfield, A.; Thorson, J.; Cohen, D. Foraging Substrate and Escape Substrate: Patch Use by Three Species of Gerbils. Ecology 2001, 82, 1781–1790, doi:10.1890/0012-9658(2001)082[1781:FSAESP]2.0.CO;2.

54. Ellard, C.G.; Eller, M.C. Spatial Cognition in the Gerbil: Computing Optimal Escape Routes from Visual Threats. Anim. Cogn. 2009, 12, 333–345, doi:10.1007/s10071-008-0193-9.

55. Lay, D.M. The Anatomy, Physiology, Functional Significance and Evolution of Specialized Hearing Organs of Gerbilline Rodents. J. Morphol. 1972, 138, 41–120, doi:10.1002/jmor.1051380103.

56. Emerson, S.E.; Kotler, B.P.; Sargunaraj, F. Foraging Efficiency in the Face of Predation Risk: A Comparative Study of Desert Rodents. Evol. Ecol. Res. 2018, 19, 61–70.

57. Panday, P.; Pal, N.; Samanta, S.; Tryjanowski, P.; Chattopadhyay, J. Dynamics of a Stage-Structured Predator-Prey Model[: Cost and Benefit of Fear-Induced Group Defense. J. Theor. Biol. 2021, 528, 110846, doi:10.1016/j.jtbi.2021.110846.

58. Kotler, B.P.; Brown, J.S.; Bouskila, A.; Mukherjee, S.; Goldberg, T. Foraging Games between Gerbils and Their Predators: Seasonal Changes in Schedules of Activity and Apprehention. Isr. J. Ecol. Evol. 2004, 50, 255–271.

59. Dall, S.R.X.; Kotler, B.P.; Bouskila, a Attention, “apprehension” and Gerbils Searching in Patches. Ann. Zool. Fennici 2001, 38, 15–23.

60. Bleicher, S.S. Multi-Predator Common Garden Experiment with Two Gerbil Species.

61. Jamovi The Jamovi Project 2023, Version 2.4.

62. Brown, J.S. Vigilance, Patch Use and Habitat Selection[: Foraging under Predation Risk. Evol. Ecol. Res. 1999, 1, 49–71.

63. Brown, J.S. Patch Use as an Indicator of Habitat Preference, Predation Risk, and Competition. Behav. Ecol. Sociobiol. 1988, 22, 37–47, doi:10.1007/BF00395696.

64. Bouskila, A. A Habitat Selection Game of Interactions between Rodents and Their Predators. Ann. Zool. Fenn. 2001, 38, 55–70.

65. Bleicher, S.S.; Kotler, B.P.; Shalev, O.; Austin, D.; Embar, K.; Brown, J.S.; Dixon, A. Divergent Behavior amid Convergent Evolution[: A Case of Four Desert Rodents Learning to Respond to Known and Novel Vipers. PLoS One 2018, 13, 1–17, doi:10.1101/362202.

66. Kotler, B.P.; Brown, J.S.; Mitchell, W.A. Environmental Factors Affecting Patch Use in Two Species of Gerbelline Rodents. J. Mammal. 1993, 74, 614–620, doi:10.2307/1382281.

67. Kotler, B.P.; Brown, J.S.; Subach, A. Temporal Foragers[: Of Optimal Coexistence of Species Mechanisms of Sand Dune Gerbils by Two Species Partitioning. Oikos 1993, 548–556.

68. Bleicher, S.S.; Ylönen, H.; Käpylä, T.; Haapakoski, M. Olfactory Cues and the Value of Information[: Voles Interpret Cues Based on Recent Predator Encounters. Behav. Ecol. Sociobiol. 2018, 72, 187–199, doi:10.1007/s00265-018-2600-9.

69. Porter, W.P.; Budaraju, S.; Stewart, W.E.; Ramankutty, N. Calculating Climate Effects on Birds and Mammals: Impacts on Biodiversity, Conservation, Population Parameters, and Global Community Structure. Am. Zool. 2000, 40, 597–630, doi:10.1093/icb/40.4.597.

70. Kotler, B.P.; Brown, J.S. Mechanisms of Coexistence of Optimal Foragers as Determinants of Local Abundances and Distributions of Desert Granivores. J. Mammal. 1999, 80, 361–374.

71. Ziv, Y.; Abramsky, Z.; Kotler, B.P.; Subach, A. Interference Competition and Temporal and Habitat Partitioning in Two Gerbil Species, 1993, Vol. 66.

72. Dixon, A.K. Tradeoffs of Food and Safety in Contrasting Environments: From the Deserts of the Mojave and the Negev to the Coral Reefs of Eilat. Ph.D. Thesis., Ben Gurion University of the Negev, 2017.

73. Blumstein, D.T.; Mari, M.; Daniel, J.C.; Ardron, J.G.; Griffin, A.S.; Evans, C.S. Olfactory Predator Recognition: Wallabies May Have to Learn to Be Wary. Anim. Conserv. 2002, 5, 87–93, doi:10.1017/S1367943002002123.

74. Thorson, J.M.; Morgan, R.A.; Brown, J.S.; Norman, J.E. Direct and Indirect Cues of Predatory Risk and Patch Use by Fox Squirrels and Thirteen-Lined Ground Squirrels. Behav. Ecol. 1998, 9, 151–157.

75. Morgan, R.A.; Brown, J.S.; Thorson, J.M. The Effect of Spatial Scale on the Functional Response of Fox Squirrels. Ecology 2013, 78, 1087–1097, doi:10.1890/0012-9658(1997)078[1087:TEOSSO]2.0.CO;2.

76. Bytheway, J.P.; Carthey, A.J.R.R.; Banks, P.B. Risk vs. Reward: How Predators and Prey Respond to Aging Olfactory Cues. Behav. Ecol. Sociobiol. 2013, 67, 715–725, doi:10.1007/s00265-013-1494-9.

77. Banks, P.B.; Bytheway, J.P.; Carthey, A.J.R.; Hughes, N.K.; Price, C.J. Olfaction and Predaotr-Prey Interactions Amongst Mammals in Australia. In Predators in Australia; Glen, A.S., Dickman, C.R., Eds.; CSIRO PUBLISHING: Collingwood, 2014; p. Chapter 17.

78. Bleicher, S.S. Heat and Humidity Alter Predation Cues in Gerbillus Andersoni Allebyi. MS Thesis. Ben Gurion University of the Negev, Ben Gurion University of the Negev, 2012.

79. Laundré, J.W.; Calderas, J.M.M.; Hernández, L. Foraging in the Landscape of Fear, the Predator’s Dilemma: Where Should I Hunt[? Open Ecol. J. 2009, 2, 1–6.

80. Vance-Chalcraft, H.D.; Soluk, D.A.; Ozburn, N. Is Prey Predation Risk Influenced More by Increasing Predator Density or Predator Species Richness in Stream Enclosures? Oecologia 2004, 139, 117–122, doi:10.1007/s00442-003-1484-0.

81. Martín, J.; López, P.; Polo, V. Temporal Patterns of Predation Risk Affect Antipredator Behaviour Allocationby Iberian Rock Lizards. Anim. Behav. 2009, 77, 1261–1266, doi:10.1016/j.anbehav.2009.02.004.

82. Abrams, P.A. Should Prey Overestimate the Risk of Predation? Am. Nat. 1994, 144, 317–328.

83. Bouskila, A.; Blumstein, D.T.; Mangel, M. Prey under Stochastic Condition Should Probably Overstimate Predation Risk: A Reply to Abrams. Am. Nat. 1995, 145, 1015–1019.

84. Bouskila, A.; Blumstein, D.T. Rules of Thumb for Predation Hazard Assessment[: Predictions from a Dynamic Model. Am. Nat. 1992, 139, 161–176.

85. Dickman, C.R. Overview of the Impacts of Feral Cats On. Nature 1996, 92.

86. Doherty, T.S.; Dickman, C.R.; Johnson, C.N.; Legge, S.M.; Ritchie, E.G.; Woinarski, J.C.Z. Impacts and Management of Feral Cats Felis Catus in Australia. Mamm. Rev. 2017, 47, 83–97, doi:10.1111/mam.12080.

87. Hardman, B.; Moro, D.; Calver, M. Direct Evidence Implicates Feral Cat Predation as the Primary Cause of Failure of a Mammal Reintroduction Programme. Ecol. Manag. Restor. 2016, 17, 152–158, doi:10.1111/emr.12210.

88. Moseby, K.E.; Peacock, D.E.; Read, J.L. Catastrophic Cat Predation: A Call for Predator Profiling in Wildlife Protection Programs. Biol. Conserv. 2015, 191, 331–340, doi:10.1016/j.biocon.2015.07.026.

89. Labra, A.; Leonard, R. Intraspecific Variation in Antipredator Responses of Three Species of Lizards (Liolaemus): Possible Effects of Human Presence. J. Herpetol. 1999, 33, 441–448.

90. Avilés-Rodríguez, K.J.; Kolbe, J.J. Escape in the City: Urbanization Alters the Escape Behavior of Anolis Lizards. Urban Ecosyst. 2019, 22, 733–742, doi:10.1007/s11252-019-00845-x.

91. Zuñiga-Palacios, J.; Zuria, I.; Castellanos, I.; Lara, C.; Sánchez-Rojas, G. What Do We Know (and Need to Know) about the Role of Urban Habitats as Ecological Traps? Systematic Review and Meta-Analysis. Sci. Total Environ. 2021, 780, 1–11, doi:10.1016/j.scitotenv.2021.146559.

92. Glen, A.S.; Dickman, C.R. Complex Interactions among Mammalian Carnivores in Australia, and Their Implications for Wildlife Management. Biol. Rev. Camb. Philos. Soc. 2005, 80, 387–401, doi:10.1017/s1464793105006718.

93. Brito, J.C.; Barrio, G. Del; Stellmes, M.; Pleguezuelos, J.M.; Saarinen, J.; Barrio, G. Del; Stellmes, M.; Pleguezuelos, J.M.; Saarinen, J. Drivers of Change and Conservation Needs for Vertebrates in Drylands[: An Assessment from Global Scale to Sahara-Sahel Wetlands an Assessment from Global Scale to Sahara-Sahel Wetlands. Eur. Zool. J. 2021, 88, 1103–1129, doi:10.1080/24750263.2021.1991496.

94. Medina, F.M.; Nogales, M. A Review on the Impacts of Feral Cats (Felis Silvestris Catus) in the Canary Islands[: Implications for the Conservation of Its Endangered Fauna. Biodivers. Conserv. 2009, 18, 829–846, doi:10.1007/s10531-008-9503-4.

95. Banks, P.B.; Dickman, C.R. Alien Predation and the Effects on Multiple Levels of Prey Naivite. Trends Ecol. Evol. 2007, 22, 229–230, doi:10.1016/j.tree.2007.02.003.

96. Carthey, A.J.R.; Banks, P.B. Naïveté in Novel Ecological Interactions: Lessons from Theory and Experimental Evidence. Biol. Rev. Camb. Philos. Soc. 2014, 89, 932–949, doi:10.1111/brv.12087.

97. Rödl, T.; Berger, S.; Romero, L.M.; Wikelski, M. Tameness and Stress Physiology in a Predator-Naive Island Species Confronted with Novel Predation Threat. Proc. Biol. Sci. 2007, 274, 577–582, doi:10.1098/rspb.2006.3755.

98. Banks, P.B.; Carthey, A.J.R.; Bytheway, J.P.; Banks, P.B.; Carthey, A.J.R.; Bytheway, J.P. Australian Native Mammals Recognize and Respond to Alien Predators□: A Meta-Analysis. 2018, 285, 1–8.

99. Ramp, D.; Benítez-lópez, A.; Carroll, S.; Carthey, A.J.R.; Rogers, E.I.E.; Zawada, K.J.A.; Middleton, O.; Lundgren, E.; Svenning, J.; Avidor, E. Savviness of Prey to Introduced Predators. 2023, 1–11, doi:10.1111/cobi.14012.

100. Serpell, J.A. Domestication and History of the Cat. In The domestic cat: The biology of Iits behavior; Turne, D.C., Bateson, P., Eds.; Cambridge Univ. Press.: Cambridge, UK, 2000; pp. 180–192 ISBN 9781139177177.

101. Banks, P.B.; Dickman, C.R.; Newsome, A.E. Ecological Costs of Feral Predator Control: Foxes and Rabbits. J. Wildl. Manage. 1998, 62, 766–772, doi:10.2307/3802353.

